# Antimicrobial resistance at the animal–environment interface: phenotypic detection and persistence of ESBL- and carbapenemase-producing Enterobacterales in Irish cattle systems

**DOI:** 10.64898/2026.09.22.752022

**Authors:** Mairead Quinn, Cormac O’Shea, Lisa Reidy

**Affiliations:** Department of Bioveterinary and Microbial Science, Technical University of the Shannon, Athlone,Ireland

**Keywords:** Antimicrobial resistance, carbapenemase-producing Enterobacterales, extended-spectrum β-lactamase, agriculture, One Health, environmental reservoir

## Abstract

Antimicrobial resistance (AMR) represents a critical One Health challenge at the interface of agriculture, the environment, and human health. In Ireland, cattle production systems generate large volumes of manure and slurry that are routinely applied to land. Yet, data on the prevalence and environmental persistence of clinically important resistance phenotypes remain limited. This study investigated the occurrence, characterisation, and persistence of extended-spectrum β-lactamase (ESBL) and carbapenemase-producing Enterobacterales (CPE), and fluoroquinolone-resistant bacteria on Irish cattle farms.

A total of 450 pooled bovine faecal samples were collected from farms across 25 counties and screened using selective culture, antimicrobial susceptibility testing, and MALDI-TOF MS identification. Carbapenemase activity in presumptive CPE isolates was confirmed using phenotypic and biochemical assays, including MALDI-TOF MBT STAR analysis to demonstrate carbapenem hydrolysis. In parallel, a controlled persistence study evaluated the survival of phenotypically resistant ESBL and CPE bacteria in bovine slurry under simulated seasonal conditions.

Overall, 29.6% of farms were positive for ESBL and/or CPE, with ESBL-only resistance most prevalent (13.7%), followed by combined ESBL–CPE phenotypes (7.5%). Fluoroquinolone resistance was frequently detected and commonly co-occurred with β-lactam resistance, indicating multidrug resistance. *Escherichia coli* was the dominant carrier of both ESBL and carbapenemase phenotypes. Persistence experiments demonstrated sustained phenotypic resistance in slurry for up to nine months under simulated winter conditions and up to three months during simulated summer conditions, with resistance becoming undetectable following extended verification.

These findings demonstrate that clinically important AMR phenotypes are embedded within Irish cattle production systems and can persist in agricultural waste streams, highlighting slurry as a significant environmental reservoir with implications for environmental dissemination and One Health surveillance.

## 1. Introduction

Antimicrobial resistance (AMR) is accelerating globally and is now recognised as one of the most urgent threats to human, animal, and environmental health. The World Health Organization (WHO) has identified AMR as a critical global challenge, associated with increased morbidity, mortality, and substantial economic burden (WHO, 2019). Within Europe, approximately one in five bacterial infections are resistant to first-line antimicrobials, with some Member States reporting resistance levels approaching 40% (EFSA, 2024). In Ireland, AMR-associated infections are estimated to cost the national health service approximately €12 million annually (Department of Health; and Department of Agriculture, 2021), while global healthcare costs attributable to AMR are projected to rise substantially by mid-century if current trends persist (ECDC, 2017, 2023; Mader *et al*., 2022).

Beyond its clinical consequences, AMR represents a complex ecological phenomenon. Resistant bacteria and resistance determinants circulate between humans, animals, and the environment, facilitated by antimicrobial use, microbial adaptation, and environmental dissemination pathways. The majority of emerging infectious diseases have zoonotic origins, and many exhibit antimicrobial resistance, underscoring the interconnected nature of these systems (Mader *et al*., 2022). Drug-resistant tuberculosis, HIV, and malaria together contribute to an estimated 700,000 deaths annually (World Health Organisation, 2023), illustrating how resistance amplifies the burden of infectious disease worldwide. These trends highlight the need for integrated surveillance and stewardship approaches grounded in a One Health framework that recognises the interdependence of human, animal, and environmental health.

Among the most significant AMR threats are extended-spectrum β-lactamase (ESBL) producing Enterobacterales. Commensal organisms such as *Escherichia coli* and *Klebsiella pneumoniae* are ubiquitous inhabitants of the gastrointestinal tract of humans and animals, yet possess a marked capacity to acquire resistance determinants and act as reservoirs for multidrug-resistant (MDR) phenotypes. ESBL-producing strains hydrolyse third-generation cephalosporins and frequently exhibit co-resistance to other critically important antimicrobial classes, including fluoroquinolones (Waade *et al*., 2021). The increasing detection of ESBL-producing Enterobacterales in both clinical and veterinary settings highlights their expanding ecological footprint and relevance across sectors (Bonnet, 2004; WHO, 2020; Tseng, Liu and Liu, 2023).

In Ireland, ESBL-producing *E. coli* accounted for 11.6% of invasive isolates in 2018, representing the highest prevalence recorded since national surveillance began (HPSC, 2020). Subsequent annual surveillance reports indicate that ESBL prevalence has remained persistently elevated, with interannual variation but no sustained decline, suggesting endemic circulation within the human population (HPSC, 2018, 2019, 2020; ECDC, 2024). High levels of combined resistance to multiple antimicrobial classes, including fluoroquinolones, have also been reported in Ireland relative to other Northern and Western European countries (ECDC, 2024). While carbapenems remain effective therapeutic options for severe ESBL associated infections in human medicine, the emergence of additional resistance mechanisms threatens to compromise their clinical utility.

Livestock are increasingly recognised as important reservoirs of ESBL-producing Enterobacterales. European surveillance data estimate the prevalence of ESBL-producing *E. coli* in food-producing animals to range between approximately 1.2% and 2.8% (ECDC, 2024); however, substantially higher prevalence has been documented at regional and herd levels. Studies from Switzerland, Germany, England, Wales, and Ireland have reported ESBL carriage in cattle populations exceeding these averages, with ESBLs detected in up to 17.2% of Irish dairy herd samples (Horton *et al*., 2011; Reist *et al*., 2013; Dahms *et al*., 2015; Ramovic *et al*., 2020). These findings raise concern regarding amplification within livestock systems and onward environmental dissemination through manure and slurry.

Carbapenemase-producing Enterobacterales (CPE) represent a further escalation in public health risk. Carbapenems are classified by the WHO as critically important antimicrobials and are widely regarded as last-resort agents for the treatment of severe MDR Gram-negative infections (WHO, 2019; Sati *et al*., 2025). In Ireland, CPE have been notifiable since 2017, reflecting their clinical significance within healthcare settings (HPSC, 2019). Although carbapenems are not authorised for veterinary use in the European Union, sporadic detections of CPE in livestock and companion animals have been reported internationally(Madec *et al*., 2017; Köck *et al*., 2018). To date, routine EU surveillance of indicator bacteria in cattle, poultry, and pigs has not detected CPE in Irish livestock (EFSA, 2024); nevertheless, their potential emergence within agricultural systems remains a critical One Health concern.

Carbapenem resistance is mediated by a diverse group of carbapenem-hydrolysing β-lactamases, most commonly encoded by bla KPC, bla NDM, bla VIM, bla IMP, and bla OXA-48 genes (Abou-assy *et al*., 2023). These enzymes belong to Ambler classes A, B, and D and differ in catalytic mechanisms and inhibitor susceptibility. Serine carbapenemases utilise an active site serine residue to cleave the β-lactam ring, whereas metallo-β-lactamases require zinc ions for hydrolytic activity (Page and Badarau, 2008). Carbapenems exert their antimicrobial effect by binding penicillin-binding proteins and disrupting peptidoglycan synthesis, leading to bacterial cell lysis (Rezaei *et al*., 2026). Increased reliance on carbapenems in human medicine has contributed to the selection and dissemination of CPE across clinical, agricultural, and environmental compartments (Meletis, 2016).

Evidence increasingly demonstrates that CPE and other carbapenem-resistant organisms are not confined to hospitals but have been detected in livestock, companion animals, wildlife, wastewater, and surface waters (Bonardi and Pitino, 2019). Environmental persistence is facilitated by co-selection with other antimicrobial classes (Selvarajan *et al*., 2023) and by plasmid-mediated horizontal gene transfer, enabling dissemination across bacterial species and ecological boundaries (Dankittipong *et al*., 2022).

Agricultural systems play a pivotal role in this ecology. Intensive livestock production has historically relied on metaphylactic and prophylactic antimicrobial use, generating selective pressure for resistant organisms within the gastrointestinal microbiome of food producing animals. Resistant bacteria may subsequently disseminate through manure and slurry, contaminating soil, water, wildlife, and crops following land application (Singh, Bhat and Ravi, 2024; Zhang *et al*., 2024). Although veterinary antimicrobial sales in the European Union have declined substantially since 2011 (EMA, 2021, 2023), fluoroquinolones remain of particular concern due to their critical importance in human medicine and their frequent association with MDR phenotypes.

Despite increasing recognition of these risks, Ireland lacks coordinated, high-resolution surveillance data describing the prevalence, distribution, and environmental persistence of ESBL-producing Enterobacterales, CPE, and other carbapenem-resistant organisms in cattle populations. In particular, the persistence of phenotypically resistant bacteria in bovine slurry under Irish climatic conditions remains poorly characterised, despite slurry representing a major pathway for environmental dissemination through land spreading and farm-to-farm transfer.

This study addresses these knowledge gaps by investigating: (i) the farm level occurrence of ESBL-producing Enterobacterales, carbapenemase-producing Enterobacterales, and fluoroquinolone-resistant bacteria in Irish cattle; (ii) the phenotypic characterisation of carbapenemase activity using complementary biochemical and MALDI–TOF based approaches; and (iii) the persistence of phenotypically resistant bacteria in bovine slurry under simulated seasonal conditions.

To our knowledge, this is the first study to combine multi-county prevalence data with functional phenotypic confirmation of carbapenemase activity and experimentally derived persistence data in bovine slurry within an Irish context. This study integrates prevalence, phenotypic characterisation, and environmental persistence to address a critical gap in Irish AMR surveillance data.

## 2. Materials and Methods

### 2.1 Study Design

This study comprised four components:

i. determination of herd-level prevalence of extended-spectrum β-lactamase (ESBL) producing Enterobacterales, carbapenemase-producing Enterobacterales (CPE), carbapenem-resistant Enterobacterales (CRE), fluoroquinolone-resistant, and multidrug resistant (MDR) isolates in Irish cattle;
ii. assessment of antimicrobial resistance among Gram-positive faecal isolates;
iii. phenotypic characterisation of carbapenemase activity in bovine CPE/CRE isolates; and
iv. evaluation of the environmental persistence of phenotypically resistant ESBL and CPE strains in bovine slurry enriched with positive control organisms.

All faecal samples were collected through the Beef Environmental Efficiency Programme – Sucklers (BEEPS-S) and the Targeted Advisory Service on Animal Health (TASAH) via Animal Health Laboratory Ireland (AHLI). No farmer-identifying information was accessed or disclosed, and no animal interventions were required for sample collection.

### 2.2 Sample Collection and Storage

A total of 450 pooled bovine faecal samples were collected from beef and dairy herds across Ireland. Each pooled sample comprised faecal material from 10 individual animals per herd. Samples were anonymised on submission, catalogued within the AHLI laboratory information management system (LIMS), and processed within seven days of collection in accordance with national guidance for carbapenemase-producing Enterobacterales (HPSC, 2018).

For long-term storage, aliquots were suspended in brain–heart infusion (BHI) broth supplemented with 15% glycerol, resulting in a final concentration of approximately 10% faecal material, and stored at −80 °C (Achá *et al*., 2005; Homeier-Bachmann *et al*., 2022). Prior to downstream analysis, samples were thawed at room temperature and homogenised using sterile swabs.

### 2.3 Isolation and Identification of Enterobacterales

Ten microlitres of homogenised faecal slurry were inoculated into 9 mL of tryptic soy broth (TSB) and incubated aerobically at 37 °C for 24 h. Enriched cultures were subsequently streaked onto CHROMagar™ ESBL and CHROMagar™ mSuperCARBA™ plates and incubated for a further 24 h at 37 °C. Colony colour and morphology were interpreted according to manufacturer instructions (CHROMagar, 2023, 2024)

Presumptive isolates were purified by quadrant streaking on tryptic soy agar and identified to species level using MALDI-TOF mass spectrometry (Biotyper, Bruker Daltonics, Germany). Only identifications achieving a log score ≥ 2.0 were accepted. Gram staining was performed in accordance with standard microbiological methods to confirm cellular morphology (Paray, Singh and Amin Mir, 2023).

### 2.4 Antimicrobial Susceptibility Testing (AST)

Antimicrobial susceptibility testing was performed using the Kirby–Bauer disk diffusion method on Mueller–Hinton agar in accordance with EUCAST interpretive criteria (EUCAST, 2017; Song, 2022). Antimicrobial disks were selected specifically to screen for ESBL production, carbapenem resistance, and fluoroquinolone resistance.

Screening for carbapenem resistance was conducted using ertapenem (10 µg) and meropenem (10 µg) disks. Isolates demonstrating reduced susceptibility to either agent were classified as presumptive CRE and subjected to further phenotypic carbapenemase testing.

Screening and confirmation of ESBL production were performed using cefotaxime (30 µg) and cefotaxime/clavulanic acid (30/10 µg) disks. Isolates exhibiting reduced susceptibility to cefotaxime were classified as presumptive ESBL producers. Phenotypic confirmation was achieved using the Combination Disk Test (CDT), whereby ESBL production was confirmed when the inhibition zone diameter around the cefotaxime/clavulanic acid disk exceeded that of cefotaxime alone by ≥ 5 mm. The Double Disk Synergy Test (DDST) was used as supportive evidence, indicated by enhancement of the inhibition zone towards the clavulanate-containing disk.

Fluoroquinolone resistance was assessed using ciprofloxacin (5 µg) disks, with resistance interpreted according to EUCAST breakpoints.

All plates were incubated at 37 °C for 18 h, and zones of inhibition were measured to the nearest millimetre.

### 2.5 Phenotypic Characterisation of Carbapenemase-Producing Enterobacterales

CPE classification in this study is based on phenotypic detection of carbapenemase activity; molecular confirmation of carbapenemase genes was not performed.

Presumptive CPE isolates were assessed using the Modified Carbapenem Inactivation Method (mCIM) and EDTA-modified CIM (eCIM) in accordance with CLSI guidelines (Dave *et al*., 2026). Reference control strains included *Klebsiella pneumoniae* (OXA-48), *Escherichia coli* (NDM-1), and *K. pneumoniae* (KPC-3).

Carbapenem hydrolysis was further evaluated using pH-based assays (litmus indicator strips and pH meter), with acidification interpreted as evidence of β-lactam hydrolysis. Functional confirmation of carbapenemase activity was performed using MALDI-TOF MBT STAR analysis (Dave *et al*., 2026), enabling detection of antibiotic degradation through mass-to-charge (m/z) shifts.

Morphological alterations following carbapenem exposure were visualised using scanning electron microscopy (SEM). Isolates were fixed, dehydrated, sputter-coated with gold, and examined using standard protocols as previously described (I. Piroeva *et al*., 2013).

### 2.6 Screening of Gram-Positive Isolates

Thirty faecal samples were randomly selected for assessment of antimicrobial resistance among Gram-positive bacteria. Samples were cultured on mannitol salt agar supplemented with either meropenem or cefotaxime/clavulanic acid. Distinct colonies were sub-cultured, purified, and identified using MALDI-TOF MS.

This screening was exploratory in nature and intended to characterise broader resistance patterns within the bovine faecal microbiota rather than to infer clinically relevant carbapenemase production.

### 2.7 Persistence Study (Phenotypic Resistance)

To evaluate environmental persistence of phenotypically resistant bacteria, bovine slurry was collected from a 47-head beef herd previously confirmed negative for ESBL and CPE carriage. Slurry aliquots were spiked with *Proteus mirabilis* (CPE) and *Klebsiella pneumoniae* ATCC 700603 (ESBL) as positive control strains.

Samples were incubated under simulated winter (4 °C) and summer (ambient temperature) conditions for a total duration of nine months. Monthly subsamples were analysed using selective CHROMagar™ media, AST, and MALDI-TOF MS to confirm species identity and resistance phenotype. Persistence was defined as continued recovery of phenotypically resistant organisms by culture and susceptibility testing, rather than genetic confirmation of resistance determinants.

### 2.8 Statistical Analysis

Statistical analyses were conducted using GraphPad Prism v8.0.2 (GraphPad Software, USA).

Agreement between phenotypic methods (chromagar) and MALDI-TOF Biotyper analysis was assessed using Cohen’s kappa (κ) and McNemar’s test.

## 3. Results

### 3.1 Farm-Level Prevalence of ESBL, CPE, and Fluoroquinolone Resistance

Of the 450 farms sampled, 133 (29.6%) yielded at least one extended-spectrum β-lactamase (ESBL)-producing or carbapenemase-producing Enterobacterales (CPE) isolate from pooled faecal samples. ESBL-only resistance was detected on 62 farms (13.7%), CPE-only resistance on 8 farms (1.8%), and concurrent ESBL–CPE resistance on 34 farms (7.5%).

Fluoroquinolone resistance was identified on 29 farms (6.4%). Exclusive fluoroquinolone resistance occurred on only 2 farms (0.5%). Co-resistance patterns included ESBL– fluoroquinolone resistance on 9 farms (2.0%) and CPE–fluoroquinolone resistance on 3 farms (0.6%). Multi-drug resistance (ESBL, CPE, and fluoroquinolone) was observed on 15 farms (3.3%).

### 3.2 Species Distribution of ESBL- and CPE-Positive Isolates

MALDI-TOF MS identification (log score ≥ 2.0) confirmed 121 ESBL-positive and 62 CPE-positive isolates. *Escherichia coli* was the predominant species, accounting for 59 of 121 ESBL isolates (48.8%) and 30 of 62 CPE isolates (48.4%). Additional taxa recovered included *Proteus* spp., *Citrobacter* spp., and *Klebsiella* spp., together with other Gram-negative organisms that grew on selective media.

Among presumptive CPE isolates, species identification by MALDI-TOF Biotyper showed substantial agreement with chromogenic carbapenemase detection (Cohen’s κ > 0.6). McNemar’s test indicated no systematic bias between the two methods (*p* > 0.05), supporting the reliability of the combined phenotypic and proteomic approach.

### 3.3 Antimicrobial-Resistant Gram-Positive Isolates

Thirty phenotypically resistant non-Enterobacterales isolates recovered from faecal samples were selected for further characterisation on the basis of growth on supplemented media and reduced antimicrobial susceptibility. These isolates were taxonomically diverse, consistent with a heterogeneous population rather than clonal expansion.

Of these, 22 were Gram-positive and belonged primarily to the families Enterococcaceae, Bacillaceae, Micrococcaceae, and Saccharomycetaceae. The most frequently identified species were *Enterococcus faecium* (*n* = 6), *Enterococcus mundtii* (*n* = 3), *Bacillus cereus* (*n* = 3), and *Bacillus licheniformis* (*n* = 3).

All presumptive carbapenem-resistant isolates were tested using the modified Carbapenem Inactivation Method (mCIM) and the EDTA-modified Carbapenem Inactivation Method (eCIM). Isolates were classified as serine carbapenemase producers (*n* = 16), metallo-β-lactamase (MBL) producers (*n* = 2), intermediate (*n* = 28), or negative (*n* = 16). Twenty isolates were advanced for confirmatory downstream analyses.

Isolates classified as intermediate produced inhibition zone diameters or minimum inhibitory concentrations that fell between the susceptible and resistant breakpoints defined by CLSI (2024). This category indicates reduced susceptibility rather than frank resistance and may still permit clinical efficacy under optimised dosing or when the drug achieves high concentrations at the infection site. It can also represent an early stage of emerging resistance.

### 3.5 Evidence of β-Lactam Hydrolysis by pH Change

All 16 serine carbapenemase-positive isolates produced significant acidification of the assay medium after carbapenem exposure, consistent with β-lactam hydrolysis (mean pH reduction 0.5 units). Phenotypically negative isolates showed no measurable pH change under the same conditions.

### 3.6 Scanning Electron Microscopy

Scanning electron microscopy of 20 isolates after carbapenem exposure revealed marked morphological alterations relative to untreated controls, including surface irregularities and structural distortions indicative of cellular damage under antibiotic pressure.

### 3.7 MALDI-TOF MBT STAR Confirmation of Carbapenemase Activity

MALDI-TOF MBT STAR assays confirmed carbapenemase-mediated β-lactam hydrolysis in 9 of the 20 isolates tested, as evidenced by characteristic mass-to-charge shifts. The remaining 11 isolates showed no detectable hydrolytic activity under the assay conditions.

### 3.8 Persistence of Phenotypically Resistant Bacteria in Bovine Slurry under Simulated Seasonal Conditions

Under simulated seasonal conditions, phenotypically resistant bacteria remained detectable in slurry for up to three months in summer and up to nine months in winter. Resistance was confirmed throughout the sampling period by selective culture and disk diffusion assays.

## 4. Discussion

### 4.1 Key findings

This study provides multi-county, farm-level evidence that clinically important antimicrobial resistance (AMR) phenotypes are present within Irish cattle production systems and that phenotypically resistant Enterobacterales can persist for prolonged periods in bovine slurry under environmentally relevant conditions. The combined use of large-scale prevalence screening, functional phenotypic confirmation of carbapenemase activity (including MALDI– TOF MBT STAR analysis), and controlled persistence experiments addresses a significant evidence gap in Ireland, where cattle production is extensive and slurry generation is substantial (Central Statistics Office, 2021).

Importantly, the detection of ESBL-producing and carbapenemase-producing Enterobacterales, frequently in combination with fluoroquinolone resistance, highlights the presence of multidrug-resistant bacteria within cattle-associated microbial communities. These findings support the view that livestock and agricultural waste streams should be considered integral components of AMR ecology, rather than isolated or secondary reservoirs (Baker *et al*., 2022; Prendergast *et al*., 2022; Silva *et al*., 2025).

### 4.2 Irish cattle systems as environmentally relevant AMR reservoirs

Ireland’s pasture-based dairy and beef sectors generate large volumes of manure and slurry, creating repeated opportunities for resistant bacteria to circulate within farms and enter surrounding ecosystems through storage and land application (Baker *et al*., 2022). While AMR surveillance in Ireland is well developed in the human clinical sector, comparatively few studies have quantified AMR carriage within the national cattle herd or examined the persistence of resistant phenotypes in slurry under seasonally relevant conditions.

These findings suggest that resistant Enterobacterales are present within Irish cattle systems and may be maintained within livestock-associated microbial communities. Slurry represents a particularly important interface between livestock and the environment, with the potential to disseminate resistant bacteria into soil and water following land application (Silva *et al*., 2025). Although direct transmission pathways to humans were not assessed in this study, the persistence of resistant phenotypes in slurry highlights a plausible mechanism for environmental dissemination.

### 4.3 ESBL carriage and fluoroquinolone co-resistance in context

ESBL-producing Enterobacterales were widely detected in cattle faecal samples in this study, indicating substantial intestinal carriage within Irish cattle populations. Earlier Irish work has reported lower ESBL prevalence in cattle; however, differences in study design, sample size, and the inclusion of environmental samples alongside faecal material complicate direct comparison. In contrast, faecal-based studies from comparable cattle production systems in Great Britain and continental Europe have reported higher ESBL carriage rates in beef and dairy herds, with prevalence estimates more closely aligned with those observed here (Velasova *et al*., 2019; Waade *et al*., 2021; Weber *et al*., 2021).

Taken together, these data suggest that ESBL carriage in Irish cattle may be underrepresented in routine surveillance outputs and that faecal sampling provides a more sensitive indicator of intestinal colonisation and potential for environmental shedding. The frequent co-occurrence of ESBL phenotypes with fluoroquinolone resistance further indicates multidrug resistance and is consistent with plasmid-mediated co-selection described in livestock-associated Enterobacterales. From a One Health perspective, this combination of resistance traits is particularly relevant due to the importance of both antimicrobial classes in human medicine.

### 4.4 Carbapenemase-producing Enterobacterales in livestock: significance without implying veterinary carbapenem use

Carbapenems are last-resort antimicrobials for the treatment of severe multidrug-resistant Gram-negative infections and are not authorised for use in food-producing animals. They are classified as critically important antimicrobials by the World Health Organisation and designated as “Category A: Avoid” for veterinary use by the European Medicines Agency, reflecting the need to preserve their effectiveness in human medicine.

Against this regulatory background, the detection of carbapenemase-producing Enterobacterales (CPE) phenotypes in cattle is of particular significance. Importantly, these findings do not imply veterinary carbapenem use. More plausible explanations include environmental contamination, indirect exposure via shared ecological reservoirs, or human-to-animal transmission pathways. Such mechanisms are consistent with broader European evidence indicating sporadic detection of CPE within the food chain and agricultural environments, despite strict restrictions on veterinary carbapenem use.

The predominance of Escherichia coli among CPE-positive isolates in this study mirrors patterns reported elsewhere and reinforces its role as a sentinel organism for monitoring the emergence and dissemination of high-risk resistance determinants across sectors.

### 4.5 Interpreting phenotypic carbapenemase confirmation: methodological value and resistance instability

A key strength of this study lies in the use of complementary phenotypic approaches to characterise carbapenemase activity in livestock-associated isolates. This was confirmed during statistical analysis, there was substantial agreement between the chromagar and MALDI-TOF Biotyper analysis using Cohen’s kappa (Cohen’s κ > 0.6) and McNemar’s test (*p* > 0.05). This validation was important as both forms of analysis were integral to the study.

Carbapenem resistance in Gram-negative bacteria can arise through multiple mechanisms, including carbapenemase production, reduced membrane permeability, and efflux pump activity. By focusing on functional assays that directly assess β-lactam hydrolysis, this study prioritised confirmation of resistance phenotypes with clear relevance to environmental persistence and transmission risk. Yet it is acknowledged that a limitation of this study is the absence of molecular characterisation of resistance genes, which would be required to confirm the genetic basis and mobility of the observed phenotypes.

Initial screening using mCIM and eCIM assays revealed heterogeneity among presumptive CPE isolates, identifying serine carbapenemase producers, metallo-β-lactamase producers, intermediate phenotypes, and isolates lacking demonstrable carbapenemase activity. Acidification following carbapenem exposure provided additional phenotypic evidence of β-lactam hydrolysis, consistent with established hydrolysis-based detection principles.

MALDI–TOF MBT STAR analysis offered a further level of functional confirmation by directly detecting hydrolysis and decarboxylation-associated mass shifts of carbapenem substrates. Concordance between mCIM/eCIM classification and MBT STAR results was high for isolates demonstrating stable carbapenemase activity, supporting the value of combining growth-based and mass spectrometry–based phenotypic methods for AMR surveillance in non-clinical settings.

Conversely, the absence of detectable hydrolytic activity in a subset of isolates and the subsequent loss of phenotypic resistance following subculture highlight the dynamic and potentially unstable nature of plasmid-mediated resistance outside of sustained antimicrobial selection pressure. This instability is particularly relevant when interpreting environmental and livestock surveillance data, where transient or inducible resistance phenotypes may be detected that do not persist over time.

### 4.6 Environmental persistence in slurry: seasonal risk and mitigation implications

The persistence experiments demonstrated that phenotypically resistant ESBL- and CPE-positive bacteria can be maintained in bovine slurry for extended periods, particularly under cooler autumn and winter conditions. Because both seasonal simulations originated from the same homogenised slurry source and temperature was the primary variable, these findings support a biologically plausible seasonal effect, whereby reduced microbial turnover at lower temperatures prolongs resistance retention.

Slurry represents a complex and dynamic microbial environment with fluctuating nutrient availability and microbial interactions that may support the survival of resistant bacteria even in the absence of direct antimicrobial exposure. The observed reduction and eventual loss of detectable resistance under warmer conditions are consistent with increased microbial competition, decay, and potential plasmid instability.

From a practical perspective, these findings support the rationale underpinning slurry storage and resting practices as a mitigation measure prior to land application. While this study does not define an optimal storage interval, it provides empirical evidence that seasonal conditions materially influence the persistence of phenotypically resistant bacteria and should be considered in environmental risk assessment and manure management strategies. Experimental conditions were designed to approximate seasonal variation, but they may not fully capture the complexity of on-farm slurry systems.

### 4.7 Gram-positive findings: environmental circulation rather than ESBL or carbapenemase biology

Carbapenem resistance remains predominantly a Gram-negative phenomenon driven by enzymatic mechanisms such as ESBLs and carbapenemases. Gram-positive bacteria do not produce these enzymes but may exhibit reduced susceptibility through alternative mechanisms, including intrinsic low-level resistance or altered penicillin-binding proteins.

In this study, the Gram-positive taxa identified, particularly Enterococcus spp. and environmental genera such as Bacillus and Micrococcus, are likely reflective of environmental exposure and the permeability of farm–environment microbial boundaries rather than clinically significant carbapenemase biology. Nevertheless, their detection reinforces the ecological framing of AMR as a system-wide phenomenon involving diverse microbial communities across animal and environmental interfaces.

### 4.8 Implications for One Health surveillance in Ireland

Data describing carbapenemase-producing Enterobacterales in food-producing animals remain limited across Europe, constraining comprehensive risk assessment and source attribution. The present findings suggest that rare but high-consequence resistance phenotypes can be detected within agricultural systems and that agricultural waste streams can sustain these phenotypes long enough to plausibly contribute to onward dissemination.

Given established links between hospital-associated CPE transmission and adjacent environmental compartments, bidirectional cycling between humans, animals, and shared environments is plausible in principle. These findings underscore the value of surveillance strategies that integrate livestock, manure and slurry, and environmental sampling alongside human clinical monitoring.

### 4.9 Limitations

This study has several limitations. First, resistance characterisation was based on phenotypic methods without molecular confirmation of resistance genes. Second, pooled sampling may underestimate within-herd variability. Third, persistence experiments were conducted under controlled conditions and may not fully reflect field environments. Despite these limitations, the study provides robust multi-county data and functional evidence of resistance persistence in an under-characterised setting.

## 5. Conclusion

This study demonstrates that clinically important antimicrobial resistance phenotypes are present within Irish cattle production systems and can persist for extended periods in agricultural waste under environmentally relevant conditions. Farm-level detection of ESBL-producing and carbapenemase-producing Enterobacterales, frequently accompanied by fluoroquinolone resistance, indicates that multidrug-resistant bacteria are embedded within livestock-associated microbial communities rather than occurring as isolated events. The predominance of Escherichia coli as a carrier species aligns with international observations and reinforces its value as a sentinel organism for resistance monitoring across sectors.

A key contribution of this work is the functional confirmation of carbapenemase activity using complementary phenotypic approaches, including MALDI–TOF MBT STAR analysis, which enabled discrimination between stable carbapenemase producers and transient or inducible resistance phenotypes. The persistence experiments further revealed that phenotypically resistant bacteria can be maintained in bovine slurry for up to nine months under cooler seasonal conditions, highlighting slurry as a significant environmental reservoir with the potential to facilitate onward dissemination.

Collectively, these findings underscore the need for integrated One Health surveillance that encompasses livestock, manure management, and environmental compartments alongside human clinical monitoring. By providing the first multi-county assessment of these resistance phenotypes in Irish cattle and novel evidence of their environmental persistence, this study offers critical data to inform antimicrobial stewardship, environmental risk assessment, and future mitigation strategies within Irish and comparable agricultural systems. Future work incorporating molecular characterisation will be essential to further resolve the genetic basis and transmission dynamics of these resistance phenotypes.

## 6. Acknowledgements

The authors thank the Animal Health Laboratory Ireland (AHLI) for providing anonymized samples and logistical support. We also acknowledge the technical assistance of [Dr. Daniel Fitzpatrick/CISD] during laboratory, SEM and MALDI–TOF analysis.

## 7. Conflict of Interest Statement

The authors declare no conflicts of interest.

## 8. Funding

This work was supported by TUS President’s Doctoral Scholarship.

---

**Figure. 3.1.**
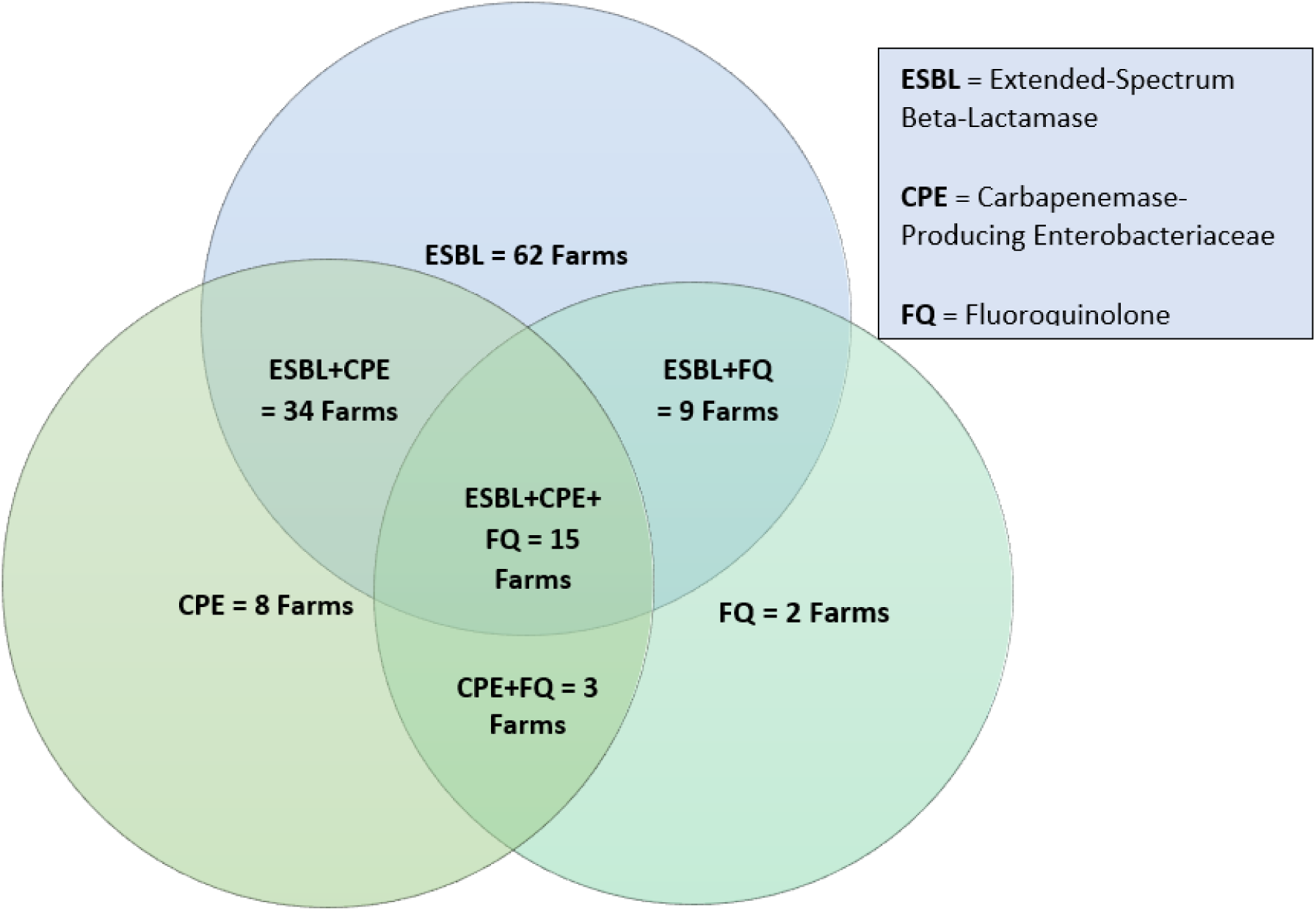
Venn diagram displaying the incidence of multi-drug resistance within the 133 AMR positive faecal samples.

**Table 3.2.** Species of bacteria recovered from bovine faecal samples and resistance demonstrated by isolates.

| Bacteria Species | ESBL | CPE |
| --- | --- | --- |
| *Escherichia coli* | 59 | 30 |
| *Proteus* sp. | 18 | 6 |
| *Citrobacter* sp. | 12 | 6 |
| *Klebsiella* sp. | 6 | 4 |
| *Achromabacter* sp. | 6 | 3 |
| *Ochrobacter* sp. | 4 | 1 |
| *Bordetella* sp. | 4 | 0 |
| *Hafnia* sp. | 3 | 2 |
| *Aliarcobacter* sp. | 3 | 0 |
| *Pseudomonas* sp. | 2 | 3 |
| *Acinetobacter* sp. | 2 | 3 |
| *Enterobacter* sp. | 1 | 1 |
| *Castellaniella* sp. | 1 | 0 |
| *Morganella* sp. | 0 | 2 |
| *Comamonas* sp. | 0 | 1 |
| **Total** | **121** | **62** |

**Figure. 3.2.1.**
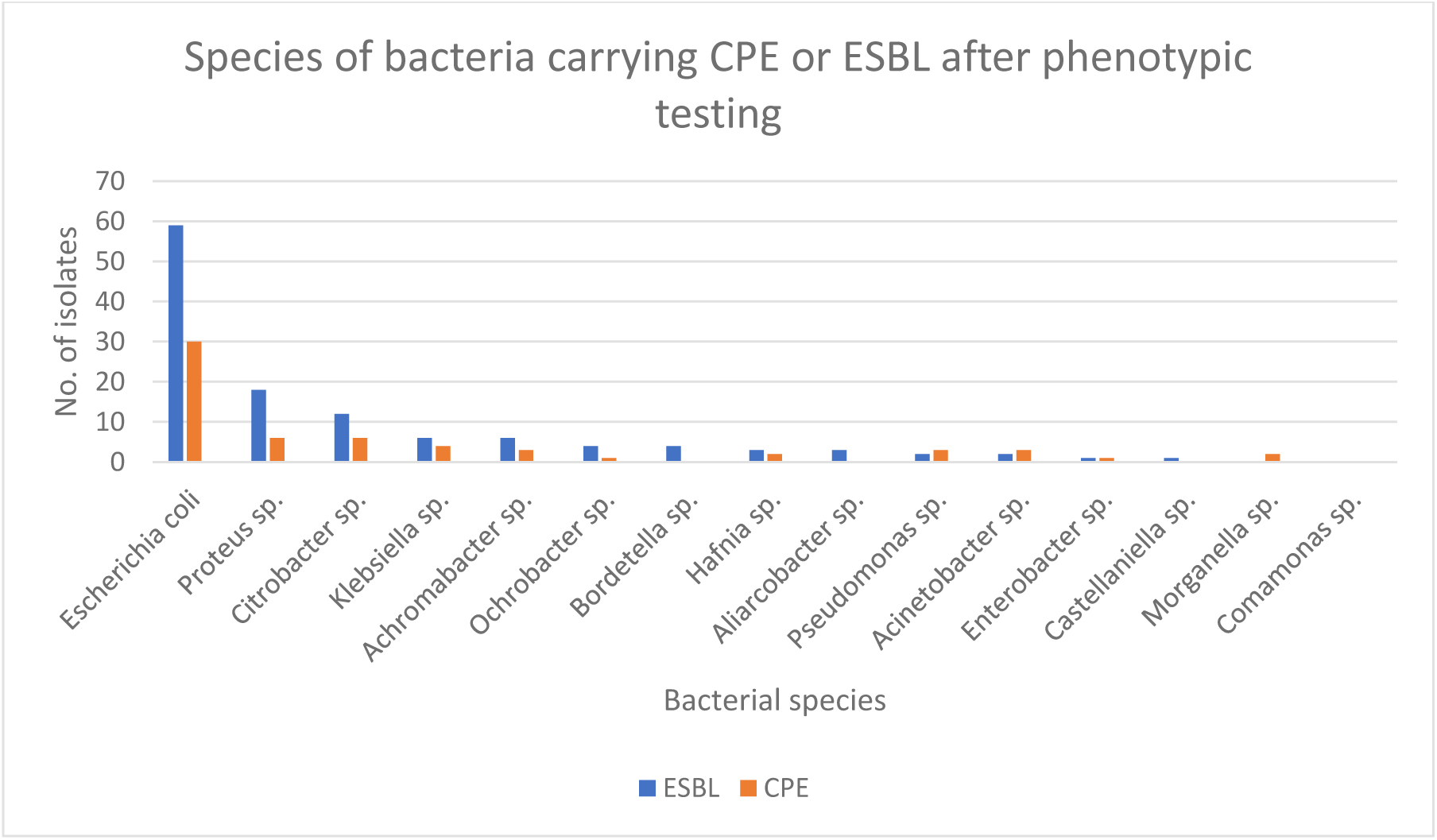
Number and species of bacteria that displayed phenotypic resistance indicating ESBL or CPE. The resistance was determined after use on selective agar and antibiotic sensitivity testing (CDT, DDST and treatment with meropenem).

**Table 3.3.** Species of non-Enterobacteriaceae detected by MALDI–TOF.

| **Bacteria** | **Family** | **Gram (+/–)** | **No. of Isolates** |
| --- | --- | --- | --- |
| *Enterococcus faecium* | Enterococcaceae | + | 6 |
| *Bacillus cereus* | Bacillaceae | + | 3 |
| *Bacillus licheniformis* | Bacillaceae | + | 3 |
| *Niallia circulans* | Bacillaceae | + | 1 |
| *Bacillus altitudinis* | Bacillaceae | + | 1 |
| *Enterococcus hirae* | Enterococcaceae | + | 2 |
| *Enterococcus faecalis* | Enterococcaceae | + | 1 |
| *Enterococcus mundtii* | Enterococcaceae | + | 3 |
| *Micrococcus luteus* | Micrococcaceae | + | 1 |
| *Debaryomyces hansenii* | Saccharomycetacea | + | 1 |

**Figure 3.4.**
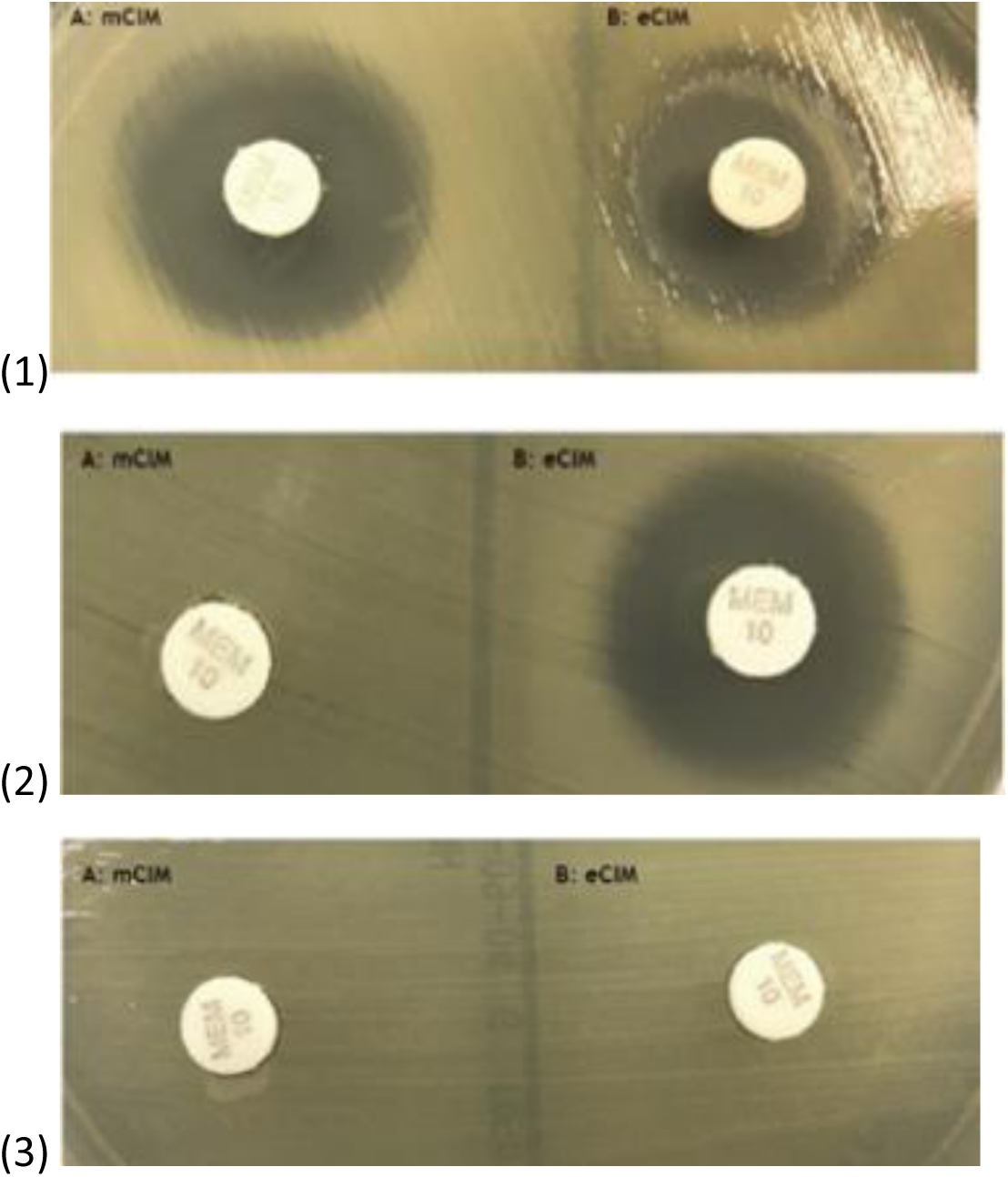
Controls for mCIM and eCIM assays: (1) CPE-negative, (2) MBL-positive, (3) Serine β-lactamase-positive.

**Table 3.5.** pH of isolates before and after meropenem treatment.

| Sample id | Bacteria species | pH of broth and bacteria solution before treatment | pH of broth and bacteria solution after treatment |
| --- | --- | --- | --- |
| 25-31-2 | *Escherichia coli* | 7.1 | 6.78 |
| 26-44-2 | *Escherichia coli* | 6.9 | 6.59 |
| 23-124-2 | *Escherichia coli* | 7.05 | 6.67 |
| 8-51-4 | *Escherichia coli* | 7.1 | 6.69 |
| 25-31-1 | *Pseudomonas stutzeri* | 7.08 | 6.5 |
| 26-44-2 | *Morganella morganii* | 7.1 | 6.7 |
| 20-44-2 | *Escherichia coli* | 7.02 | 6.59 |
| 8-51-4 | *Morganella morganii* | 6.8 | 6.4 |
| 17-94 | *Escherichia coli* | 7 | 6.6 |
| 26-6 | *Acinetobacter calcoaceticus* | 7.12 | 6.59 |
| 2-60-1 | *Escherichia coli* | 6.87 | 6.51 |
| 26-6-1 | *Escherichia coli* | 7 | 6.7 |
| 8-51-3 | *Pseudomonas stutzeri* | 7.1 | 6.83 |
| 26-49-2 | *Proteus mirabilis* | 7.05 | 6.72 |
| 23-124-2 | *Proteus mirabilis* | 7.1 | 6.71 |
| 27-5-1 | *Escherichia coli* | 6.8 | 6.56 |
| 8-53-1 | *Klebsiella pneumonia* | 7 | 7 |
| 24-143 | *Proteus mirabilis* | 7 | 7 |
| 24-115 | *Pseudomonas lundensis* | 7 | 7 |
| 25-70-1 | *Escherichia coli* | 7 | 7 |

**Figure 3.6.**
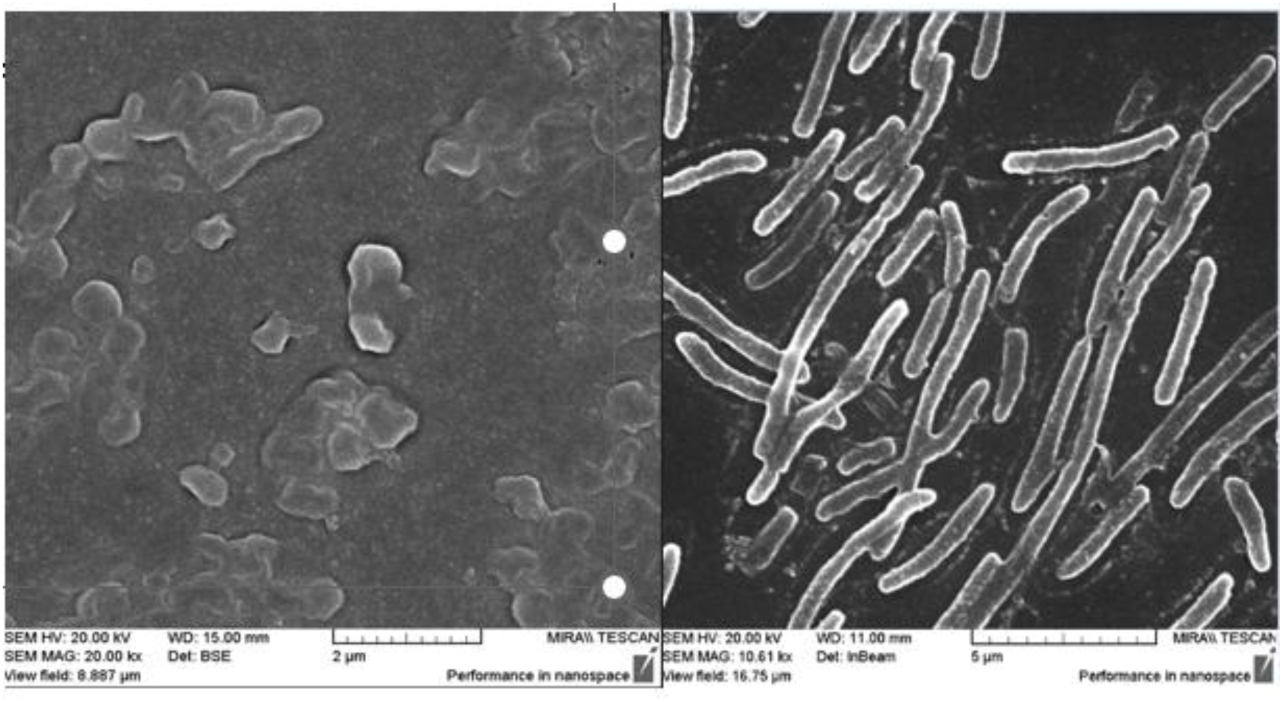
(Left) Meropenem-treated CPE-negative E. coli displaying the effects of the antibiotic on the cell integrity, structural distortion (Right) Meropenem-treated CPE-negative *E. coli which displays normal morphology*.

**Figures 3.7.**
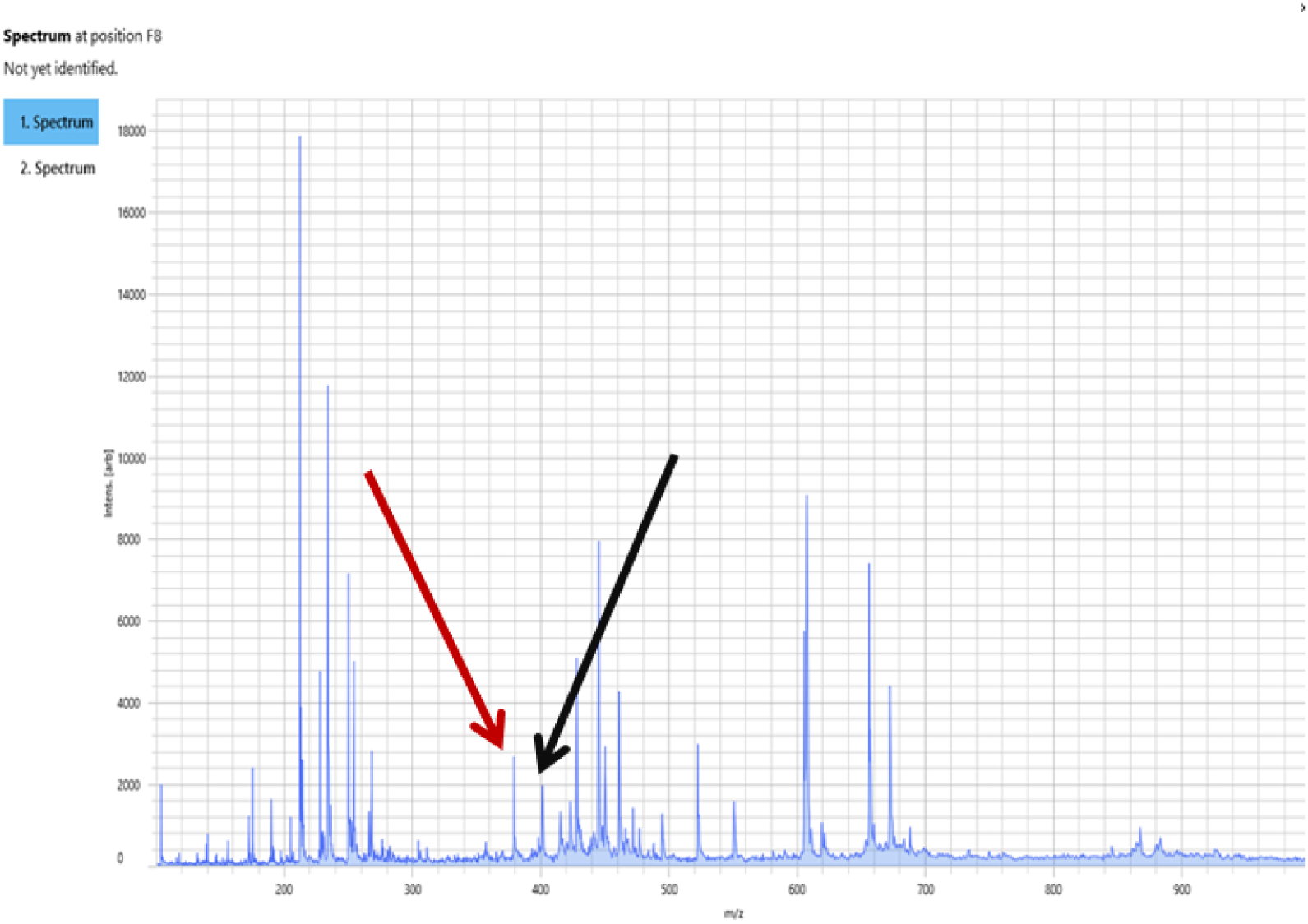
CPE positive spectra showing hydrolysed and unhydrolyzed peaks. These peaks demonstrate the mode of action of carbapenemase. This spectra is typical of a CPE positive isolate. The red arrow denotes the unhydrolyzed meropenem peak and the black arrow denotes the hydrolysed meropenem peak.

**Figures 3.7.1.**
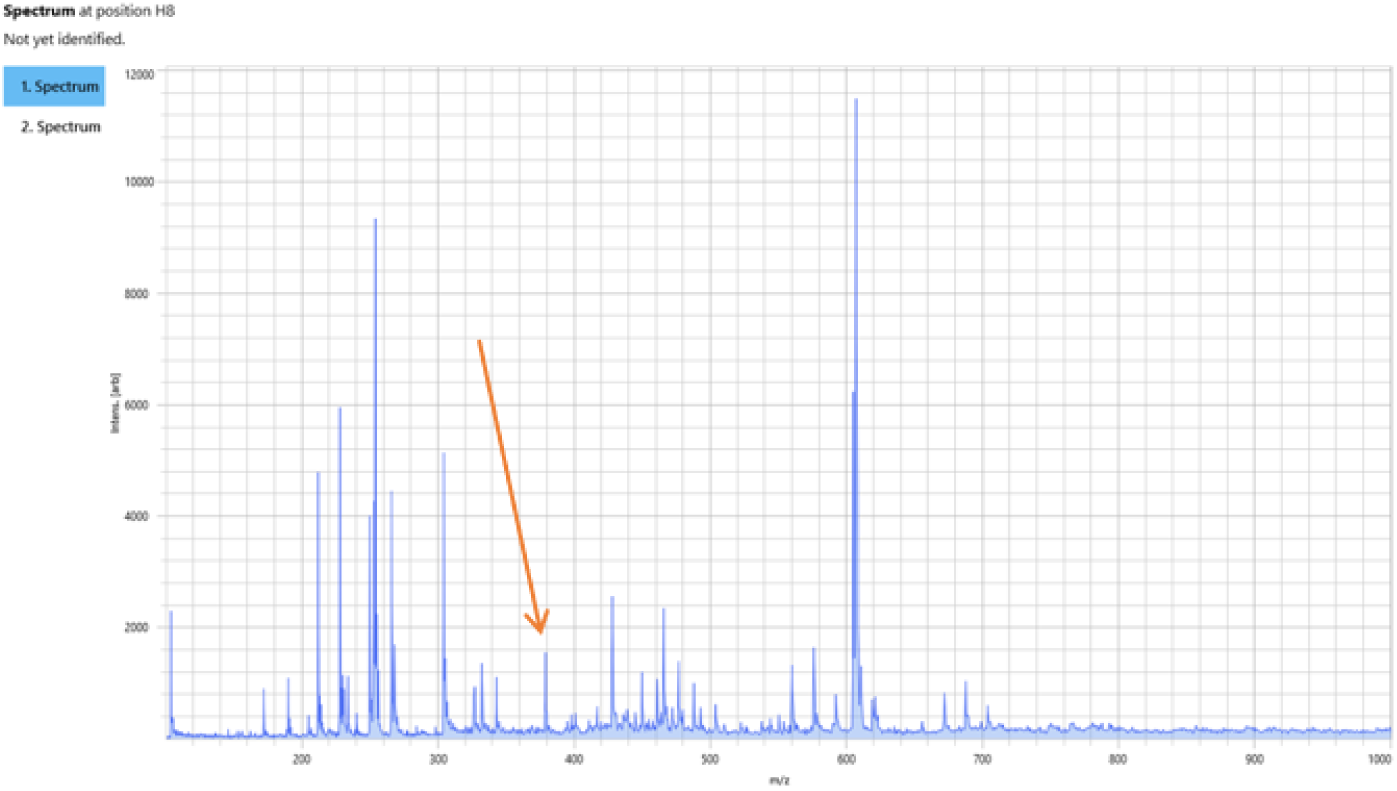
CPE negative spectra showing hydrolysed peaks. This peak demonstrates the mode of action of carbapenems. This spectra is typical of a CPE negative isolate.

**Figure. 3.8.**
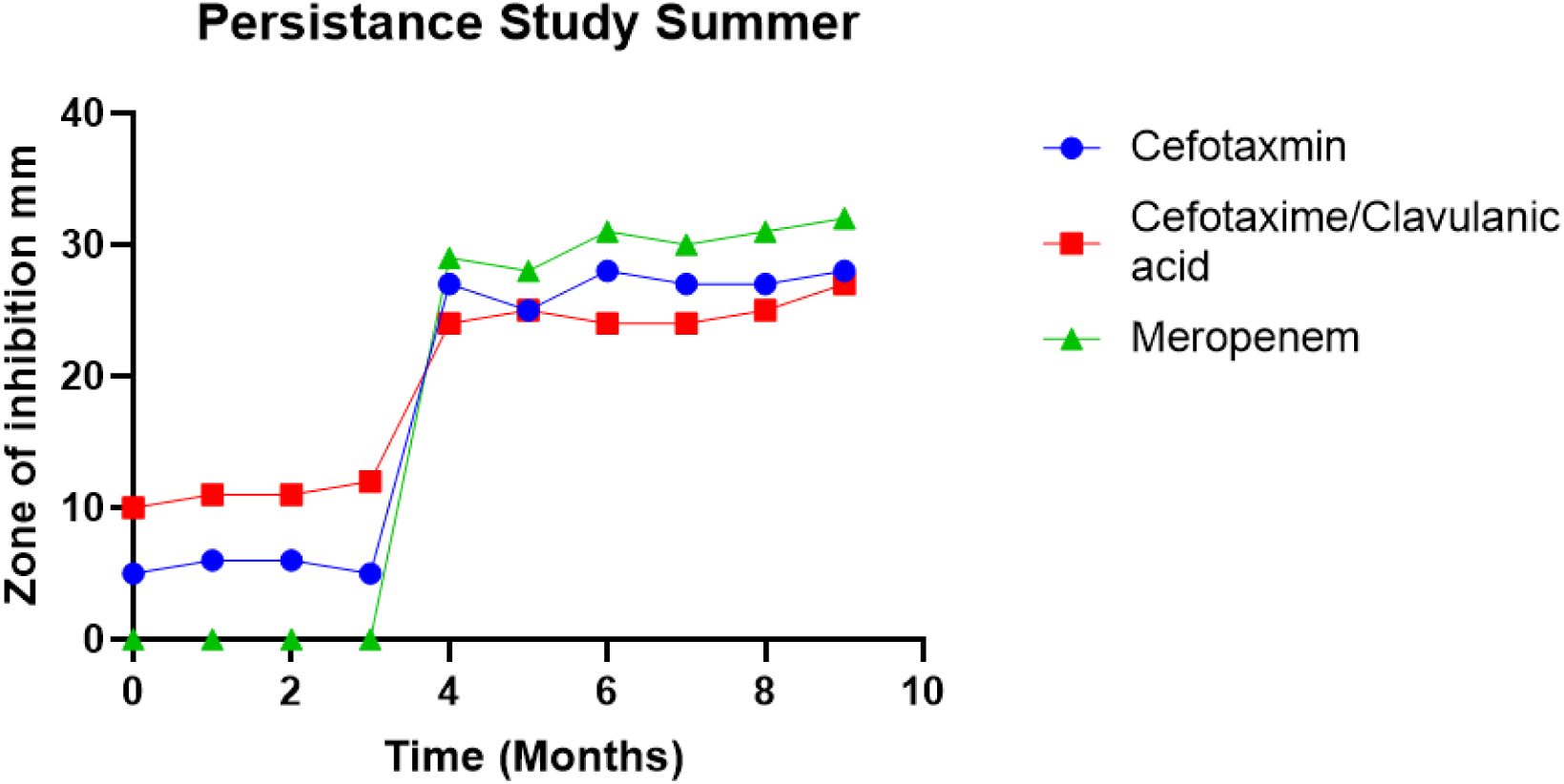
Resistance profile of summer slurry samples for both ESBL and CPE.

**Figure. 3.8.1.**
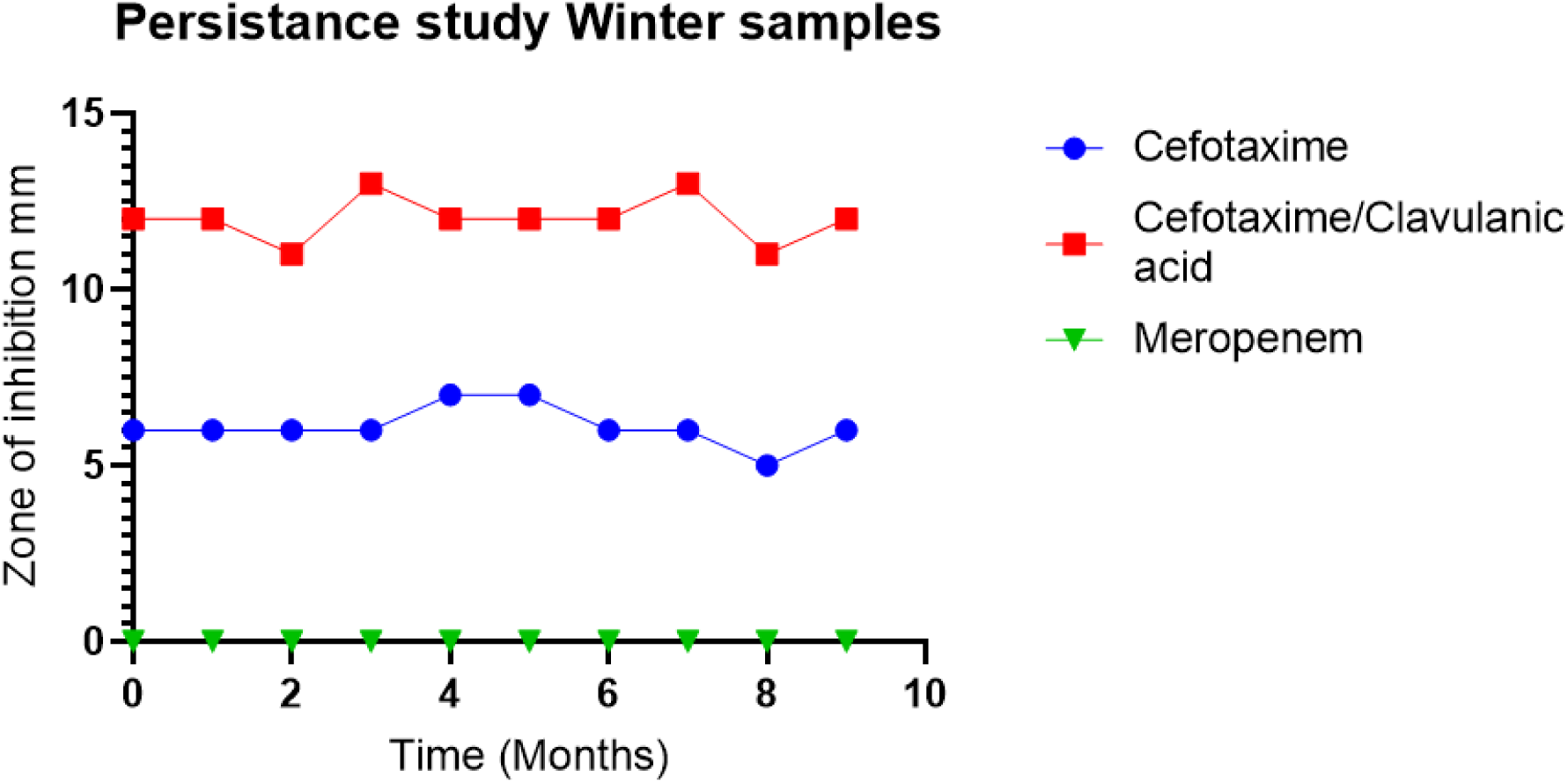
Resistance profile of winter slurry samples for both ESBL and CPE.

## Notes

### Competing Interest Statement

The authors have declared no competing interest.

